# Generation of bright autobioluminescent bacteria by chromosomal integration of the improved *lux* operon *ilux2*

**DOI:** 10.1101/2021.12.10.472100

**Authors:** Carola Gregor

**Affiliations:** Max Planck Institute for Biophysical Chemistry, Department of NanoBiophotonics, Göttingen, Germany; Institut für Nanophotonik Göttingen e.V., Department of Optical Nanoscopy, Göttingen, Germany; Cluster of Excellence “Multiscale Bioimaging: from Molecular Machines to Networks of Excitable Cells” (MBExC), University of Göttingen, Göttingen, Germany

## Abstract

The bacterial bioluminescence system enables light production in living cells without an external luciferin. Due to its relatively low levels of light emission, many applications of bioluminescence imaging would benefit from an increase in brightness of this system. In this report a new approach of mutagenesis and screening of the involved proteins is described that is based on the identification of mutants with improved properties under rate-limiting reaction conditions. Multiple rounds of screening in *Escherichia coli* resulted in the operon *ilux2* that contains 26 new mutations in the fatty acid reductase complex which provides the aldehyde substrate for the bioluminescence reaction. Chromosomal integration of *ilux2* yielded an autonomously bioluminescent *E. coli* strain with 7-fold increased brightness compared to the previously described *ilux* operon. The *ilux2* strain produces sufficient signal for the robust detection of individual cells and enables highly sensitive long-term imaging of bacterial propagation without a selection marker.

## Introduction

Cellular light emission by the process of bioluminescence can be conveniently used to image living cells. Due to its independence from external light, bioluminescence can be easily imaged with a low-complexity optical system while background signal is virtually absent. Therefore, bioluminescence is in particular superior to fluorescence imaging in samples with high autofluorescence such as tissues. In addition, bioluminescence imaging can be used for long-term measurements without phototoxicity or photobleaching and also enables the observation of light-sensitive cells.

The bioluminescence system from bacteria is of particular interest since it allows imaging without external substrates. This is due to the fact that both the light-producing enzyme, the luciferase, and its substrates can be synthesized by the cell itself by simultaneous expression of the following genes: *luxA* and *luxB* encoding the subunits of the luciferase, *luxC, luxD* and *luxE* encoding the subunits of the fatty acid reductase complex and *frp* encoding a flavin reductase. The fatty acid reductase complex produces the aldehyde substrate for the luciferase reaction from cellular fatty acids whereas the flavin reductase generates the second substrate FMNH_2_. Notably, the bacterial bioluminescence system is also functional in non-bacterial cell types such as yeast and mammalian cells ^1-4^.

One specific application of the bacterial bioluminescence system is the highly sensitive detection of bacteria in different environments. For example, it can be used to image the spreading of bacterial infections within the living host ^5-8^ or to detect bacterial contaminations in food samples ^9-11^. Since these measurements need to be performed over long time periods with quantifiable signal, continuous and constant expression of the involved genes must be ensured without the need for antibiotic selection. Therefore, it is desirable to integrate the genes into the bacterial chromosome rather than to express them from a plasmid which can be lost under non-selective conditions ^12-14^. Since in the case of chromosomal expression only one copy of the transgene is present per cell, the bioluminescence emission is generally lower than from a plasmid where the copy number is usually much higher (up to 700 copies per chromosome ^15^). Thus, it is particularly important to use protein variants of the bacterial bioluminescence system that produce high light levels in order to enable the reliable detection of even low cell numbers. In this paper, the generation of a new improved *lux* operon called *ilux2* is described that can be used for chromosomal labeling of bacteria with enhanced light emission, allowing even the detection of single *E. coli* cells.

## Results

### Design of a new screening system

Since numerous applications of bioluminescence imaging would benefit from a genetically encodable bioluminescence system with high light output, we previously optimized the brightness obtained with the *luxCDABE* operon from *Photorhabdus luminescens* in *Escherichia coli* ^16^. We observed that bioluminescence emission in *E. coli* is not only limited by the activity of the luciferase itself, but also by cellular production of the substrates (FMNH_2_ and fatty aldehyde). Consequently, incorporation of a flavin reductase (*frp*) into the *lux* operon and improving substrate production by several rounds of error-prone mutagenesis of *luxCDE* and *frp* substantially contributed to the increased light emission of the final improved operon *ilux*.

In our previous approach, we expressed all *lux* genes from a single vector pGEX(-) ^16^ at relatively high levels during error-prone mutagenesis and screening. By mutagenesis of the individual genes in *ilux* pGEX(-), it was not possible to identify any additional mutations that further increased the bioluminescence emission. Although this could indicate that the Lux proteins are already optimized, two different explanations for this observation exist: First, supply of ATP and NADPH that are required for recycling of the substrates of the bioluminescence reaction might be limited by the cellular metabolism so that no further increase in light emission can be achieved by modifications of the Lux proteins. However, this explanation can be excluded since bioluminescence was increased by expressing *ilux* at higher levels from the vector pQE(-) instead of pGEX(-) ^16^. Second, it is possible that further mutations in *ilux* which enhance the brightness exist, but that their effect is too small to detect them under the screening conditions used, although several such mutations combined might still be able to increase the bioluminescence emission substantially.

To explore this second possibility further, a mathematical model of the bioluminescence reaction was developed in order to determine the screening conditions under which the screening is most sensitive (i.e., even small changes in enzyme activities can be detected). Since the only observable in the screening is the bioluminescence signal rather than the enzyme activities themselves, this implies that the relative increase of the bioluminescence signal with an increase in activity of the iLux proteins should be maximized. For this purpose, the rate of light emission B as a function of the substrate concentrations was first qualitatively calculated by assuming the reaction scheme in Figure 1 ^17-25^ with all rate constants set to 1 for simplicity, as explained in the *Materials and Methods* section (*F*: FMNH_2_, *A*: aldehyde, *O*: molecular oxygen, *L*: luciferase):

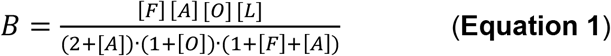

**Figure 1.**
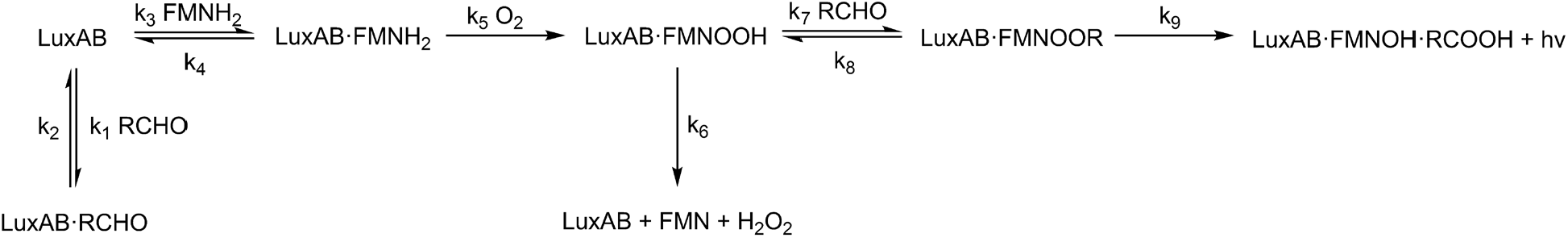
Reaction scheme of the bacterial bioluminescence reaction. LuxAB: luciferase, RCHO: fatty aldehyde, RCOOH: fatty acid, FMN: flavin mononucleotide, FMNH_2_: reduced FMN, O_2_: molecular oxygen, H_2_O_2_: hydrogen peroxide, FMNOOH: C4a-peroxy-FMN, FMNOOR: FMN-C4a-peroxyhemiacetal, FMNOH: C4a-hydroxy-FMN.

As shown in Figure 2A, the bioluminescence signal first increases with the FMNH_2_ and aldehyde concentrations and decreases again at high aldehyde levels as a consequence of aldehyde inhibition resulting from reversible formation of a luciferase-aldehyde complex ^22^, with the optimal aldehyde concentration depending on the FMNH_2_ concentration. As can be seen from Equation 1, bioluminescence increases less than proportionally with the substrate concentrations. Therefore, an improvement in activity of the substrate-producing enzymes is not reflected to the same extent in the bioluminescence signal, and hence even significantly improved variants may not be detectable in the screening. Under ideal screening conditions, the relative increase in enzymatic activity would closely approach the increase in bioluminescence. To determine the optimal conditions for screening, the relative change of the bioluminescence signal with the concentrations of FMNH_2_ and aldehyde was therefore calculated (see *Materials and Methods* section). As can be seen from Figure 2C–E, the relative change in signal is highest when production of the substrate of interest is low and therefore rate-limiting for the overall bioluminescence process, whereas the other substrate is present in excess. Likewise, the luciferase is assumed to be rate-limiting for the bioluminescence reaction if high concentrations of both FMNH_2_ and aldehyde are present. To implement this in the new screening system, the components LuxAB, LuxCDE and Frp were analyzed separately. The component to be improved by mutagenesis was expressed at low levels from a pGEX(-) vector containing a lac promoter and medium-copy-number p15A origin of replication (ori) whereas the other components were expressed at high levels from a separate plasmid using the stronger tac promoter and high-copy-number ColE1 ori (Table 1).

**Figure 2.**
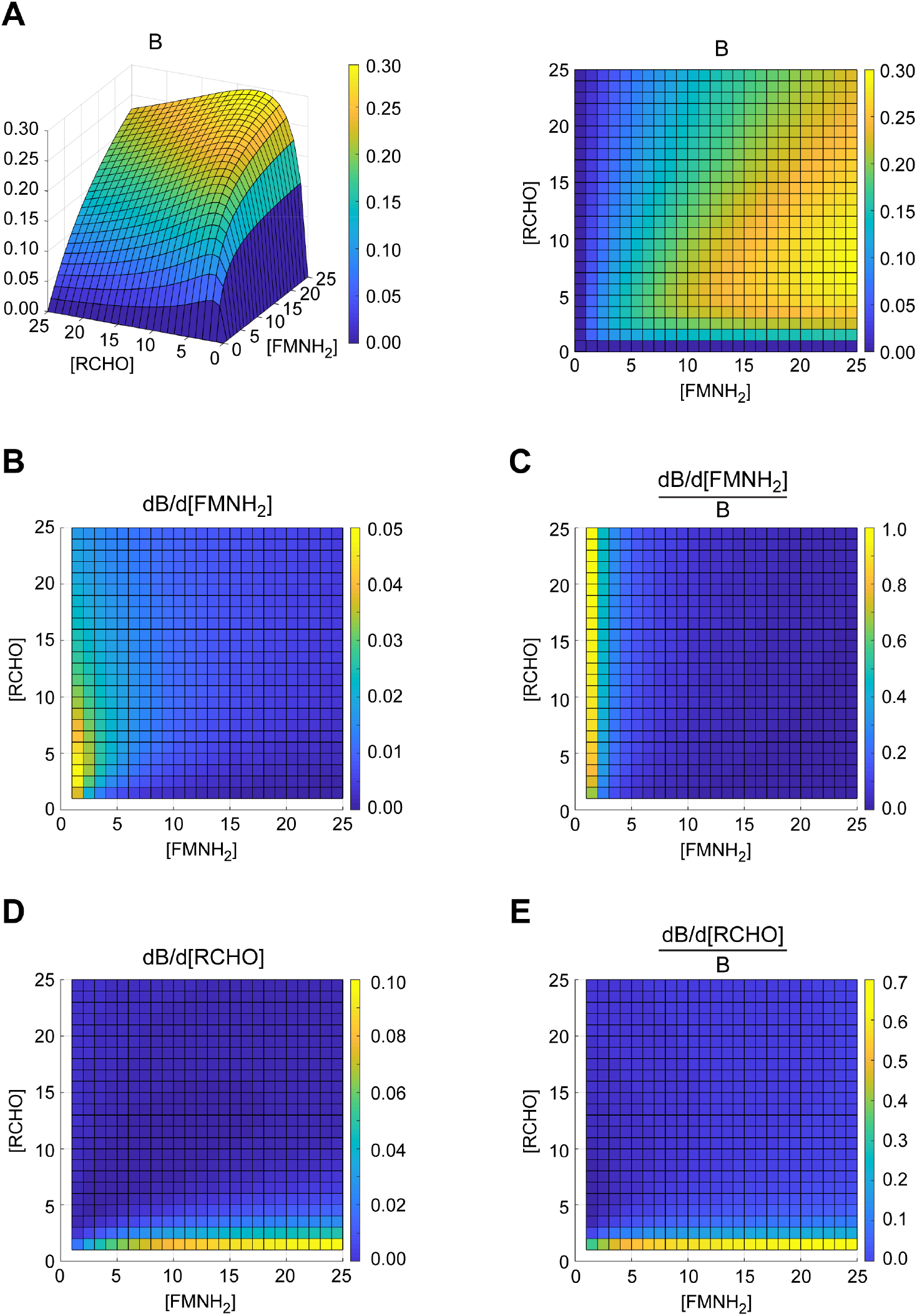
Qualitative model of the bioluminescence signal. (**A**) Bioluminescence signal B as a function of the concentrations of the substrates RCHO (aldehyde) and FMNH_2_. B was calculated based on the reaction scheme in Fig. 1 by setting all rate constants and the concentrations of luciferase and oxygen to 1. Left: side view, right: top view. (**B**) Derivative of the bioluminescence signal B with respect to the concentration of FMNH_2_, plotted as a function of the substrate concentrations. (**C**) Derivative of B with respect to the concentration of FMNH_2_, normalized to B. (**D**) Derivative of B with respect to the concentration of RCHO. (**E**) Derivative of B with respect to the concentration of RCHO, normalized to B. High values in (C) and (E) represent large relative changes of the bioluminescence signal with the respective substrate concentration and therefore optimal conditions for screening. Values are represented by their color as indicated in the colorbar in arbitrary units.

**Table 1.** Plasmid combinations used for error-prone mutagenesis. pGEX(-) was used as the vector backbone.

| Plasmid for error-prone mutagenesis |  |  |  | Plasmid containing the remaining *lux* genes |  |  |  |
| --- | --- | --- | --- | --- | --- | --- | --- |
| Insert | Promoter | Origin of replication | Resistance | Insert | Promoter | Origin of replication | Resistance |
| *iluxAB* | lac | p15Aori | ampicillin | *iluxCDEfrp* | tac | ColE1 | kanamycin |
| *iluxCDE* | lac | p15Aori | ampicillin | *iluxABfrp* | tac | ColE1 | kanamycin |
| *ilux frp* | tac | p15Aori | ampicillin | *iluxCDABE* | tac | ColE1 | kanamycin |

### Generation of *ilux2*

Starting from the previously described *ilux* operon ^16^, the three components *iluxAB, iluxCDE* and *ilux frp* were modified by repeated rounds of error-prone mutagenesis in independent screenings. Mutagenesis of *iluxAB* resulted in the new variant *iluxXAB* that contains 18 new mutations (Table 2) and produced fivefold higher bioluminescence signal under screening conditions than *iluxAB* (Figure 3A). However, when expressed at higher levels together with *iluxCDEfrp* from the same vector, *iluxXAB* yielded substantially lower signal than *iluxAB* (Figure 3B,C) and inhibited cell growth at high expression levels for unknown reasons. *iluxXAB* was therefore not further investigated.

**Table 2.** Mutations identified during the screening of *luxAB* in addition to the mutations in *ilux*.

| Gene | Mutations |
| --- | --- |
| *iluxXA* | L24M, I76V, Q86H, M139L, K283R, N289S, I298T, Y360- |
| *iluxXB* | T28S, Q95H, D112G, K193R, K238N, Y276F, M306T, V311A, D317A, H323L |

**Figure 3.**
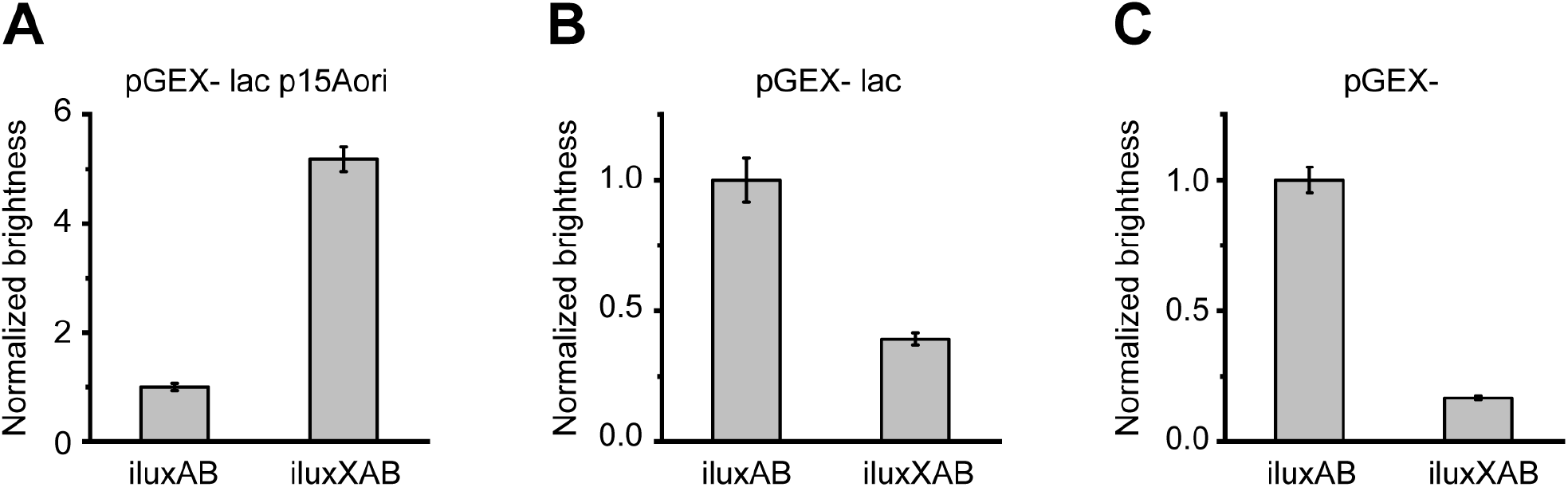
Mutagenesis and screening of *iluxAB*. Plots show the relative bioluminescence signal normalized to *iluxAB*. (**A**) For mutagenesis of *iluxAB, iluxAB* was expressed at low levels from the vector pGEX(-) lac p15Aori whereas *iluxCDEfrp* was simultaneously expressed at high levels from pGEX(-) Kan. Repeated rounds of mutagenesis resulted in the brighter variant *iluxXAB*. (**B**) Relative brightness of *iluxAB* and *iluxXAB* expressed from pGEX(-) lac. *iluxCDEfrp* was expressed from the same vector. (**C**) Relative brightness of *iluxAB* and *iluxXAB* expressed from pGEX(-). *iluxCDEfrp* was expressed from the same vector. Error bars represent SD of 5 different clones.

Mutagenesis and screening of *iluxCDE* yielded the improved variant *ilux2CDE* containing 26 new mutations in the *luxC* and *luxD* genes (Table 3). Under screening conditions, *ilux2CDE* exhibited a 160-fold higher brightness than *iluxCDE* (Figure 4A). When expressed at increasing levels from the same plasmid as *iluxABfrp* (Figure 4B–D), the difference in brightness decreased as expected (Figure 4B,C) since LuxCDE activity became less limiting for the bioluminescence process. At very high expression levels, however, the brightness of *ilux2CDE* was decreased in comparison to *ilux* (Figure 4D). Since cell growth was concomitantly reduced, this is attributed to toxic effects resulting from excessive aldehyde production, although additionally aldehyde inhibition of the luciferase cannot be ruled out. The beneficial properties of *ilux2CDE* therefore only prevail if its expression levels are not too high.

**Table 3.** Mutations contained in *ilux2* in addition to the mutations in *ilux*.

| Gene | Mutations |
| --- | --- |
| *ilux2C* | T10N, N27T, N45T, I47T, E54D, D74N, S77P, L132I, D135G, T216I, Y248N, K345N, V348A, K414E, T445K |
| *ilux2D* | N3K, I90T, K106N, N108S, M112V, N165K, D168V, K234R, V251A, D274N, E300V |

**Table 4.**
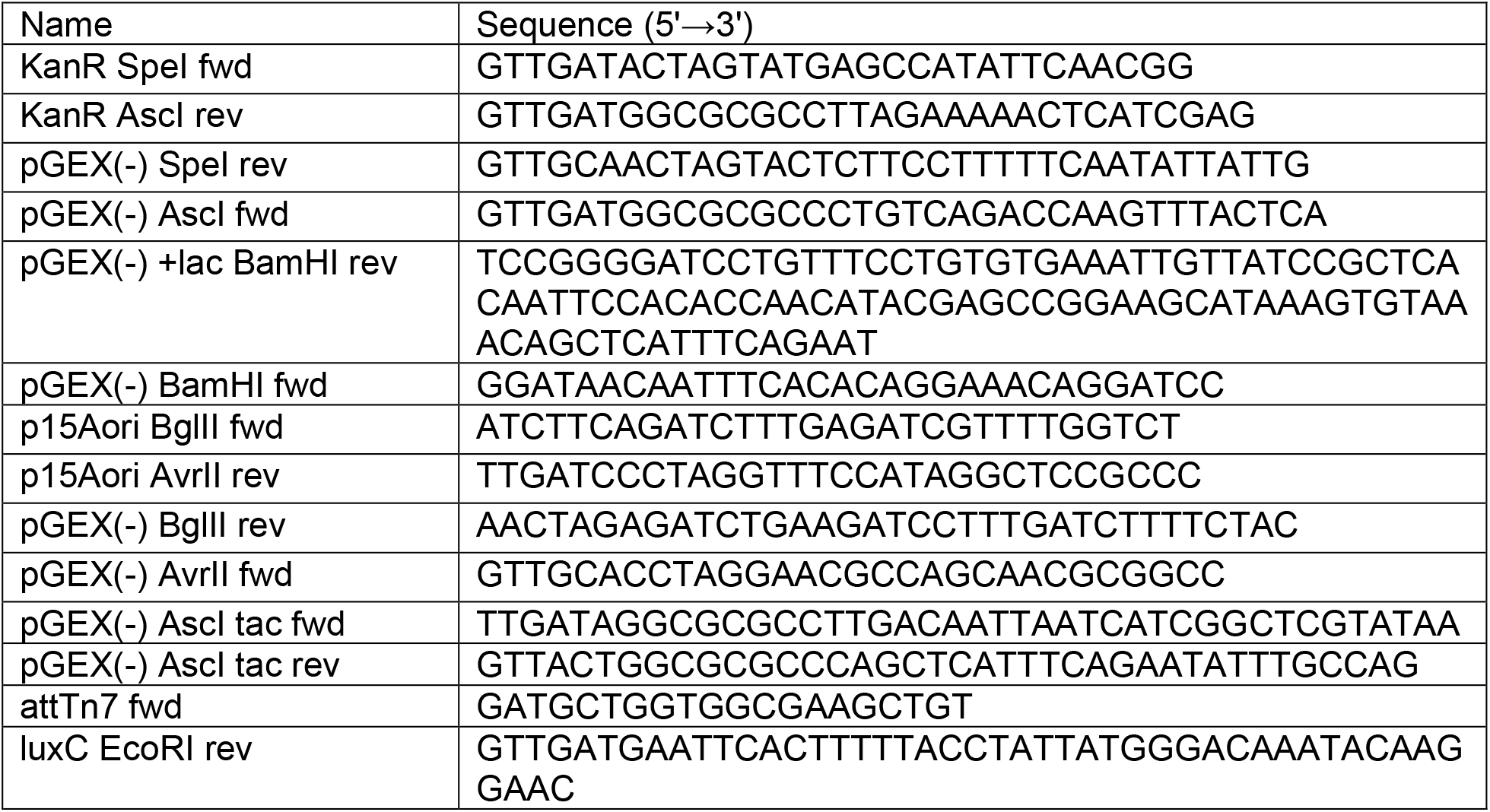
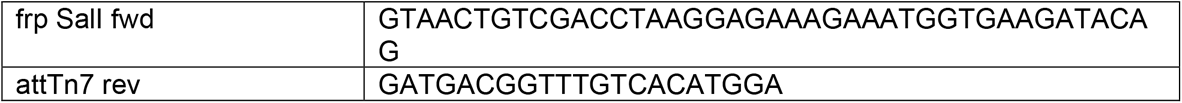
Primer sequences.

**Figure 4.**
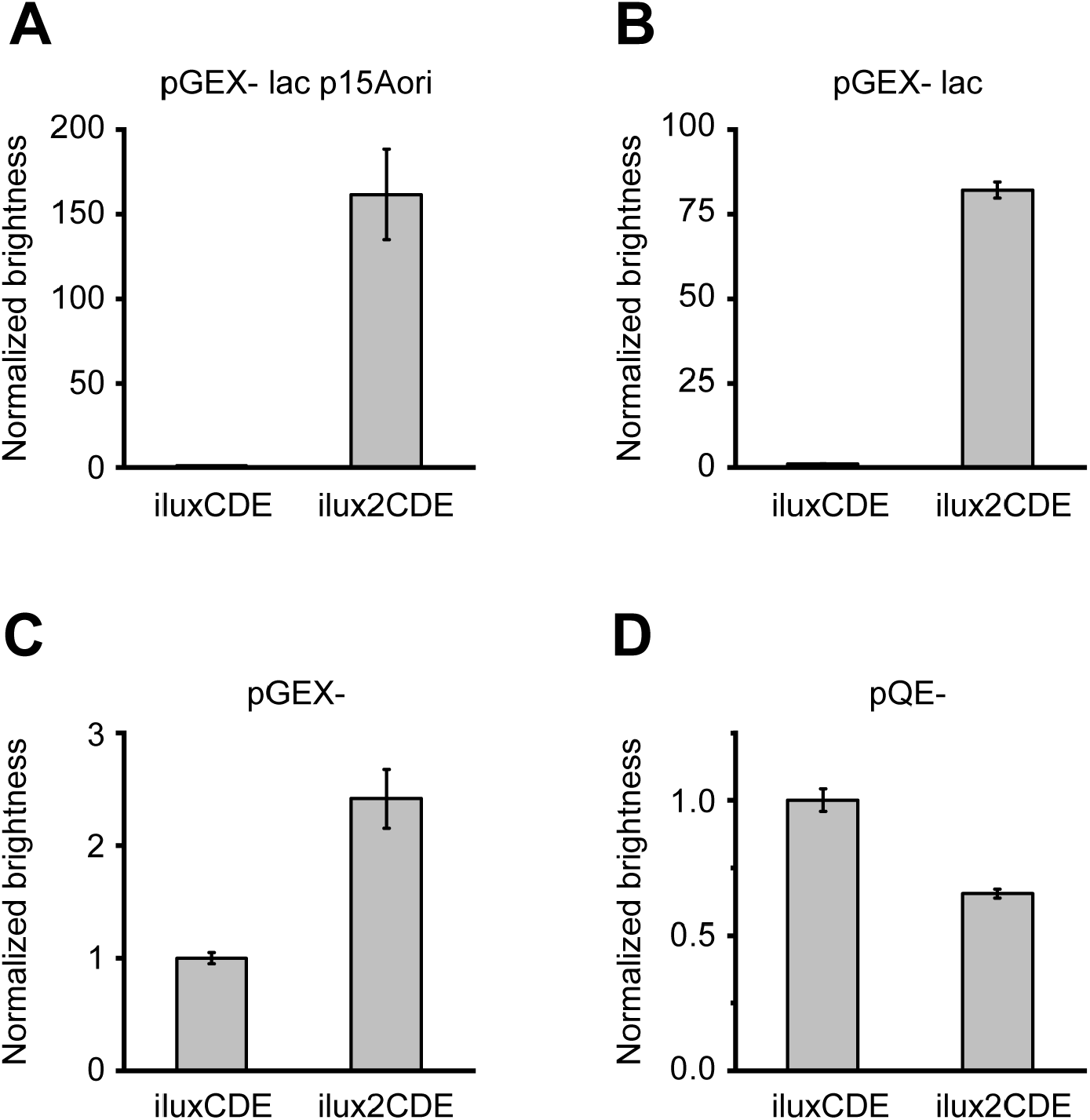
Mutagenesis and screening of *iluxCDE*. Plots show the relative bioluminescence signal normalized to *iluxCDE*. (**A**) For mutagenesis of *iluxCDE, iluxCDE* was expressed at low levels from the vector pGEX(-) lac p15Aori whereas *iluxABfrp* was simultaneously expressed at high levels from pGEX(-) Kan. Repeated rounds of mutagenesis resulted in the brighter variant *ilux2CDE*. (**B**) Relative brightness of *iluxCDE* and *ilux2CDE* expressed from pGEX(-) lac. *iluxABfrp* was expressed from the same vector. (**C**) Relative brightness of *iluxCDE* and *ilux2CDE* expressed from pGEX(-). *iluxABfrp* was expressed from the same vector. (**D**) Relative brightness of *iluxCDE* and *ilux2CDE* expressed from pQE(-). *iluxABfrp* was expressed from the same vector. Error bars represent SD of 5 different clones.

Screening of the third component *frp* required special consideration. Whereas Frp activity needed to be low in comparison to LuxAB and LuxCDE for a screening that was sensitive by the above considerations, it simultaneously had to be sufficiently high to be distinguishable from the activity of other cellular flavin reductases. Using the same plasmid combinations for screening as for *luxAB* and *luxCDE*, co-expression of *ilux frp* did not result in higher bioluminescence than expression of *iluxCDABE* alone (Figure 5A). To reduce the background signal resulting from cellular flavin reductases, an *E. coli* strain was therefore engineered in which the flavin reductase gene *fre* was knocked out since Fre has been described to be the major FMN reductase for bioluminescence in *E. coli* ^26^. However, bioluminescence from *iluxCDABE* in this strain was still not different between *ilux frp* pGEX(-) lac p15Aori and the empty vector (Figure 5B), indicating that the activity of other cellular flavin reductases was still too high. As a solution to this problem, *ilux frp* expression was increased by exchanging the lac promoter in the screening vector against the stronger tac promoter. The resulting bioluminescence could now clearly be discriminated from the empty vector (Figure 5C). Importantly, the brightness was still significantly lower than with the higher *ilux frp* expression from pGEX(-) Kan (Figure 5C) so that improvements in Frp could still increase the bioluminescence signal. However, it was not possible to identify brighter mutants by error-prone mutagenesis and screening, possibly because *ilux frp* cannot be further improved.

**Figure 5.**
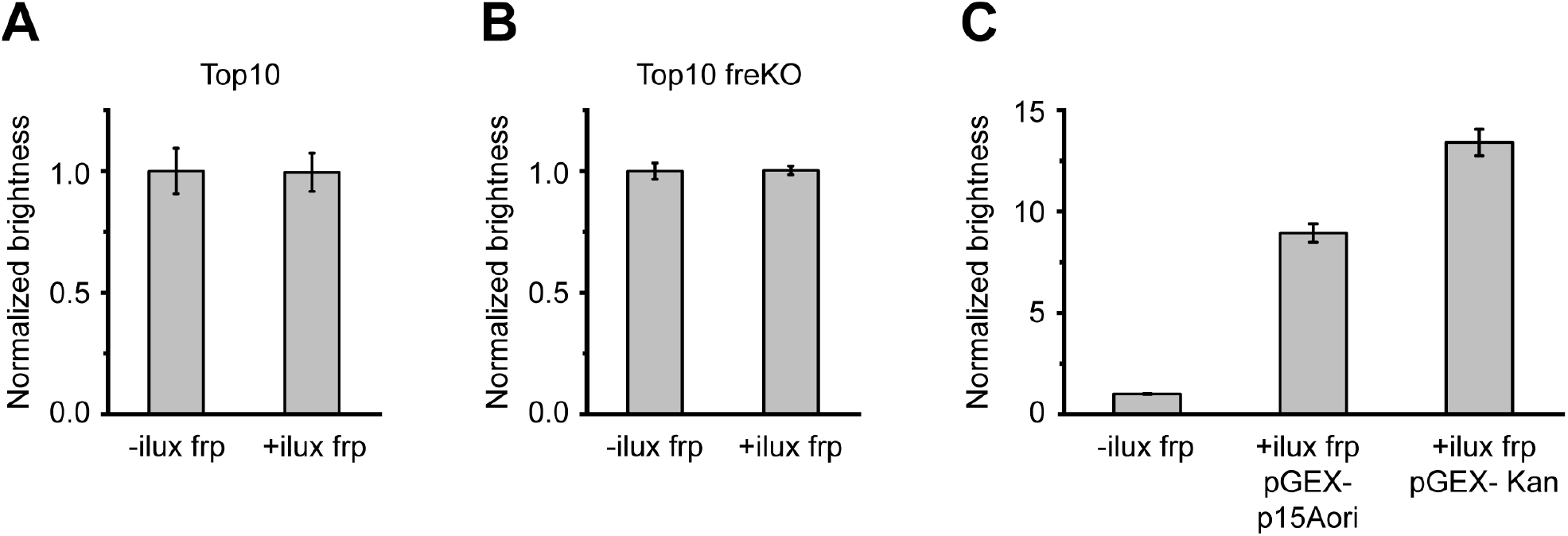
Mutagenesis and screening of *ilux frp*. Plots show the relative bioluminescence signal normalized to *iluxCDABE* without *frp*. (**A**) Relative brightness of *iluxCDABE* expressed from pGEX(-) Kan with the empty vector pGEX(-) lac p15Aori (left) and with additional expression of *ilux frp* from pGEX(-) lac p15Aori (right) in Top10 cells. (**B**) Relative brightness of *iluxCDABE* expressed from pGEX(-) Kan with the empty vector pGEX(-) lac p15Aori (left) and with additional expression of *ilux frp* from pGEX(-) lac p15Aori (right) in a Top10 *fre* knockout (freKO) strain. (**C**) Relative brightness of *iluxCDABE* pGEX(-) Kan in the Top10 freKO strain without additional expression of *ilux frp* (i.e., with the empty vector pGEX(-) p15Aori) (left), with additional expression of *ilux frp* from pGEX(-) p15Aori (middle) and with additional expression of *ilux frp* from the same vector pGEX(-) Kan (right). For mutagenesis of *ilux frp*, expression from pGEX(-) p15Aori (together with *iluxCDABE* from pGEX(-) Kan) was chosen due to the sufficiently increased brightness compared to cellular FMNH_2_ production only. Error bars represent SD of 5 different clones.

Overall, the brightness of *ilux* could only be improved by the mutations in the fatty acid reductase complex. The resulting operon was named *ilux2* which consists of *ilux2CDE* and *iluxABfrp* arranged in the original order, i.e., *ilux2CD iluxABEfrp*.

### Characterization of *ilux2*

To analyze potential changes in aldehyde chain length specificity of iLux2CDE in comparison to iLuxCDE, it was attempted to purify the LuxCDE proteins to measure their activity with different fatty acids *in vitro*. According to our previous work ^16^, no protein tags should be added to the LuxCDE proteins for this purpose since they may strongly reduce enzymatic activity of the fatty acid reductase complex. Unfortunately, purification of the untagged LuxCDE proteins in sufficient amounts in their active form was not successful. Therefore, the substrate specificity of iLuxCDE and iLux2CDE was compared under cellular conditions instead. For this purpose, *luxCDE wt, iluxCDE* and *ilux2CDE* were expressed from pGEX(-) Kan in *E. coli* and the cells were pre-incubated with saturated fatty acids of different chain lengths. The produced aldehyde was detected by adding a separate population of *E. coli* cells expressing *iluxABfrp* to the mixture. Their cell permeability allowed the aldehydes to diffuse from the *luxCDE*-into the *iluxABfrp*-expressing cells where they served as substrates for the bioluminescence reaction so that aldehyde production could be compared by the bioluminescence intensity.

Octanoic (C8), hexadecanoic (C16) and octadecanoic acid (C18) did not produce signal discriminable from the background signal caused by production of aldehydes from cellular substrates. For octanoic acid, this is likely due to the fact that aldehydes of eight or fewer carbon atoms are poor substrates for bacterial luciferase ^27, 28^, whereas in case of hexadecanoic and octadecanoic acid this may be due to the slow cellular entry of long-chain aldehydes ^29^. For decanoic (C10), dodecanoic (C12) and tetradecanoic acid (C14), the bioluminescence obtained with LuxCDE wt, iLuxCDE and iLux2CDE was compared (Figure 6). Due to potential differences in cellular uptake, the signal cannot be compared between different fatty acids, but only between the LuxCDE variants. The signal from tetradecanoic acid whose corresponding aldehyde is regarded as the natural substrate for the bacterial bioluminescence reaction ^30^ was not significantly different between LuxCDE wt, iLuxCDE and iLux2CDE. For decanoic and dodecanoic acid, iLux2CDE exhibited increased signal, whereas the brightness of iLuxCDE was surprisingly somewhat reduced compared to LuxCDE wt. It is therefore assumed that the increased bioluminescence observed with iLux2CDE is at least in part due to the enhanced utilization of cellular fatty acids other than tetradecanoic acid.

**Figure 6.**
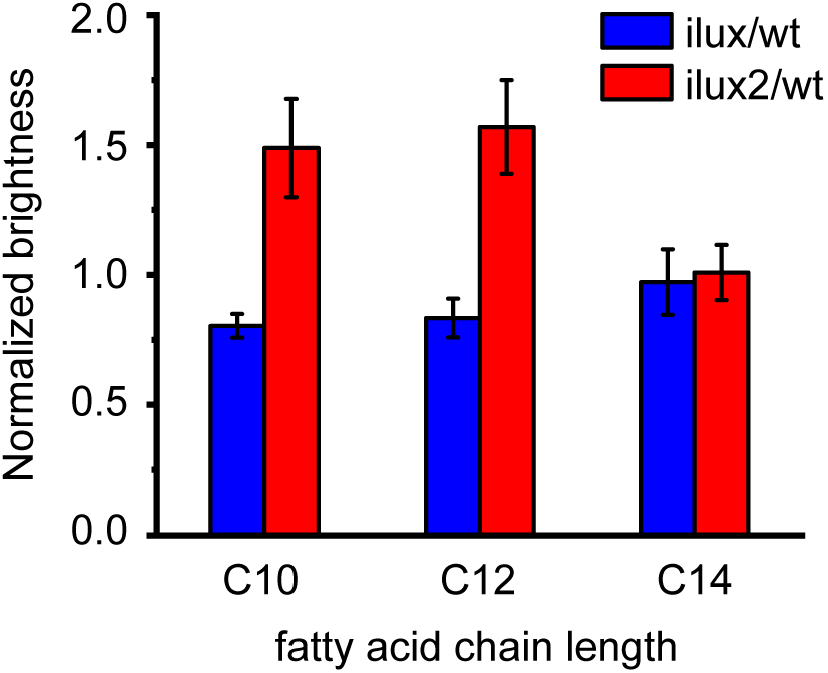
Comparison of aldehyde production of LuxCDE wt, iLuxCDE and iLux2CDE from saturated fatty acids of different chain length. Top10 cells expressing *luxCDE wt, iluxCDE* or *ilux2CDE* from pGEX(-) Kan were incubated in LB medium containing 300 µM decanoic, dodecanoic or tetradecanoic acid for 5 min. Subsequently, Top10 cells expressing *iluxABfrp* from pGEX(-) Kan were added to detect the aldehyde produced by LuxCDE by bioluminescence. Imaging was performed at room temperature. For each fatty acid, the bioluminescence signal was normalized to LuxCDE wt. Error bars represent SD of 4 different clones.

To test if iLux2CDE increases the brightness also in bacteria other than *E. coli, luxCDABE wt, ilux* and *ilux2* were expressed from the vector pJOE7771.1 ^31^ in *Pseudomonas fluorescens* (Figure 7). Both *ilux* and *ilux2* exhibited higher brightness than *luxCDABE wt* with bioluminescence emission from *ilux2* being highest, indicating that *ilux2* is also a superior reporter in bacteria other than *E. coli*.

**Figure 7.**
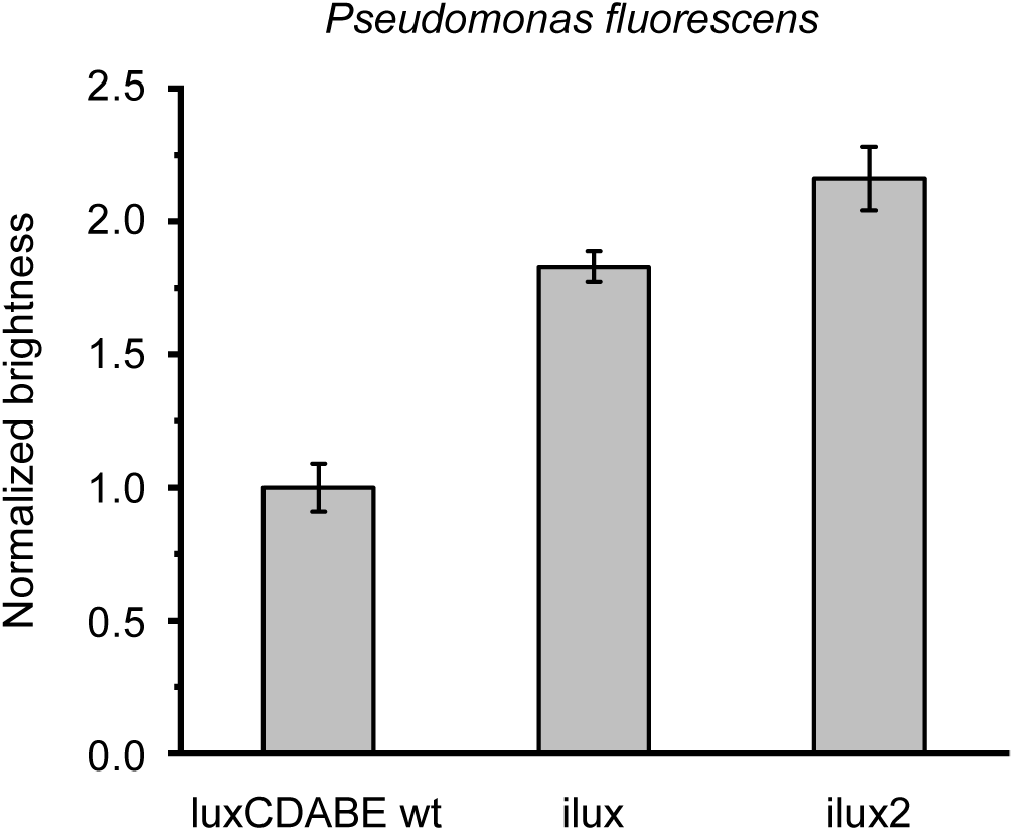
Comparison of *luxCDABE wt, ilux* and *ilux2* in *Pseudomonas fluorescens*. The indicated *lux* operons were expressed in *P. fluorescens* from the vector pJOE7771.1 at 30 °C. The bioluminescence signal was normalized to the *luxCDABE wt* operon. Error bars represent SD of 5 different clones.

Next, *E. coli* strains stably expressing the different *lux* operons were engineered by inserting them into the chromosome. Apart from the *lux* variant, the brightness also strongly depends on its expression levels, and therefore higher bioluminescence can generally be achieved with expression from a plasmid due to the higher copy number per cell. However, stable maintenance of a plasmid usually requires selection with an antibiotic resistance marker, which is not feasible or desirable for many applications such as long-term imaging of bacterial infections in living animals and plants, and hence chromosomal expression is preferable. The pGRG25 vector ^32^ was therefore used for site-specific chromosomal insertion of the *lux* operons at the *attTn7* site in *E. coli* Top10 cells (Figure 8). In the chromosomally labeled strains, bioluminescence from *ilux* was only ∼65% higher than from *luxCDABE wt*. One reason for this may be that FMNH_2_ production by other cellular enzymes is comparatively high at the lower chromosomal expression levels, and therefore its supply is less rate-limiting for the bioluminescence reaction. The brightness of the *ilux2* strain was ∼12-fold higher than that of the *luxCDABE wt* strain (∼7-fold higher than *ilux*). Compared to the highest levels of light emission that were obtained so far with *ilux* expressed from the vector pQE(-), the stable *ilux2* strain exhibits 12% of the brightness of this extrachromosomal high-copy-number system. Therefore, the *ilux2* strain should enable highly sensitive and quantifiable detection of *E. coli* cells during long-term imaging without selection pressure.

**Figure 8.**
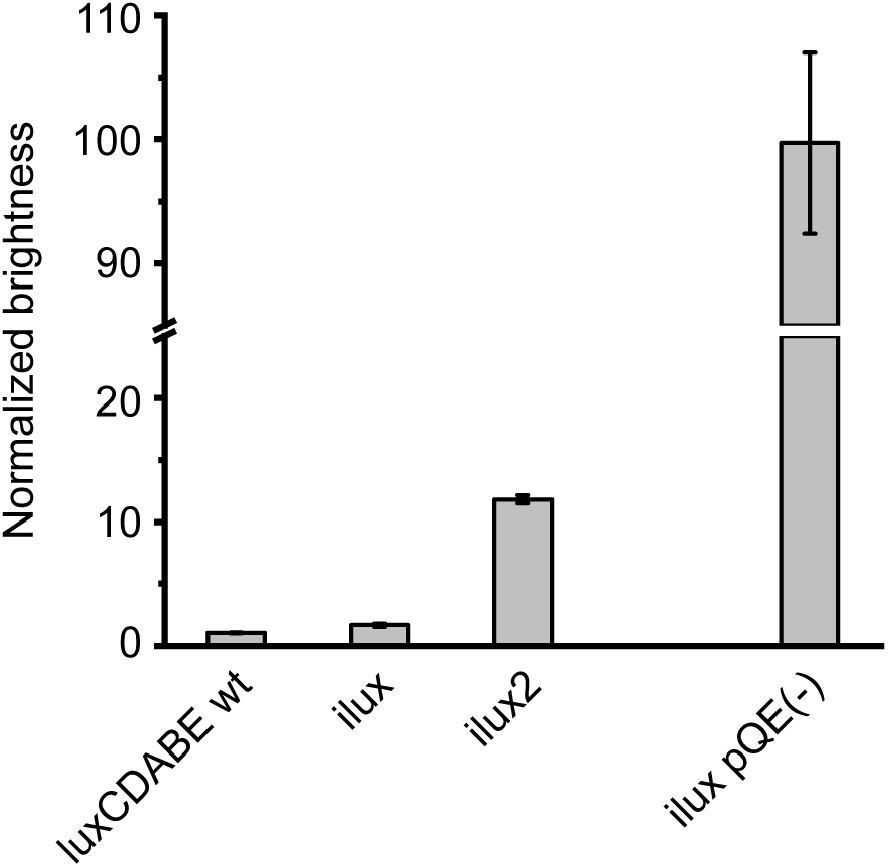
Comparison of bioluminescence emission from the chromosomally expressed operons *luxCDABE wt, ilux* and *ilux2* (left) with *ilux* expressed from the vector pQE(-) (right) in Top10 cells. The bioluminescence signal was normalized to the *luxCDABE wt* strain. Error bars represent SD of 5 different clones.

### Bioluminescence imaging of *ilux2*

The *ilux2* strain was next applied for bioluminescence imaging in different samples. First, the cell number required for detection in macroscopic samples using a commercial imaging system was determined. For this purpose, different dilutions of the stably bioluminescent *E. coli* strains were applied onto mashed potatoes and imaged with an Amersham Imager (AI) 600 RGB (Figure 9). Addition of 1 µl of cell suspension resulted in spreading of the cells on the surface of the sample within an area of ∼5 mm in diameter. At the resulting cell density, around 1·10^4^ and 7·10^3^ cells were required to reliably detect the *luxCDABE wt* and *ilux* strains, respectively, whereas only 1·10^3^ cells were required for the *ilux2* strain. In contrast, a strain chromosomally labeled with EYFP could not be detected by fluorescence with the AI 600 RGB even at the highest cell number of 10^6^ cells used. While bioluminescence imaging is limited by the low number of emitted photons, fluorescence imaging of samples with high autofluorescence such as food products or living tissues mostly suffers from the high fluorescence background. Consequently, discrimination of the fluorescence signal of the EYFP strain was impossible despite the higher brightness of the image (Figure 9D), demonstrating the superiority of bioluminescence imaging in these samples.

**Figure 9.**
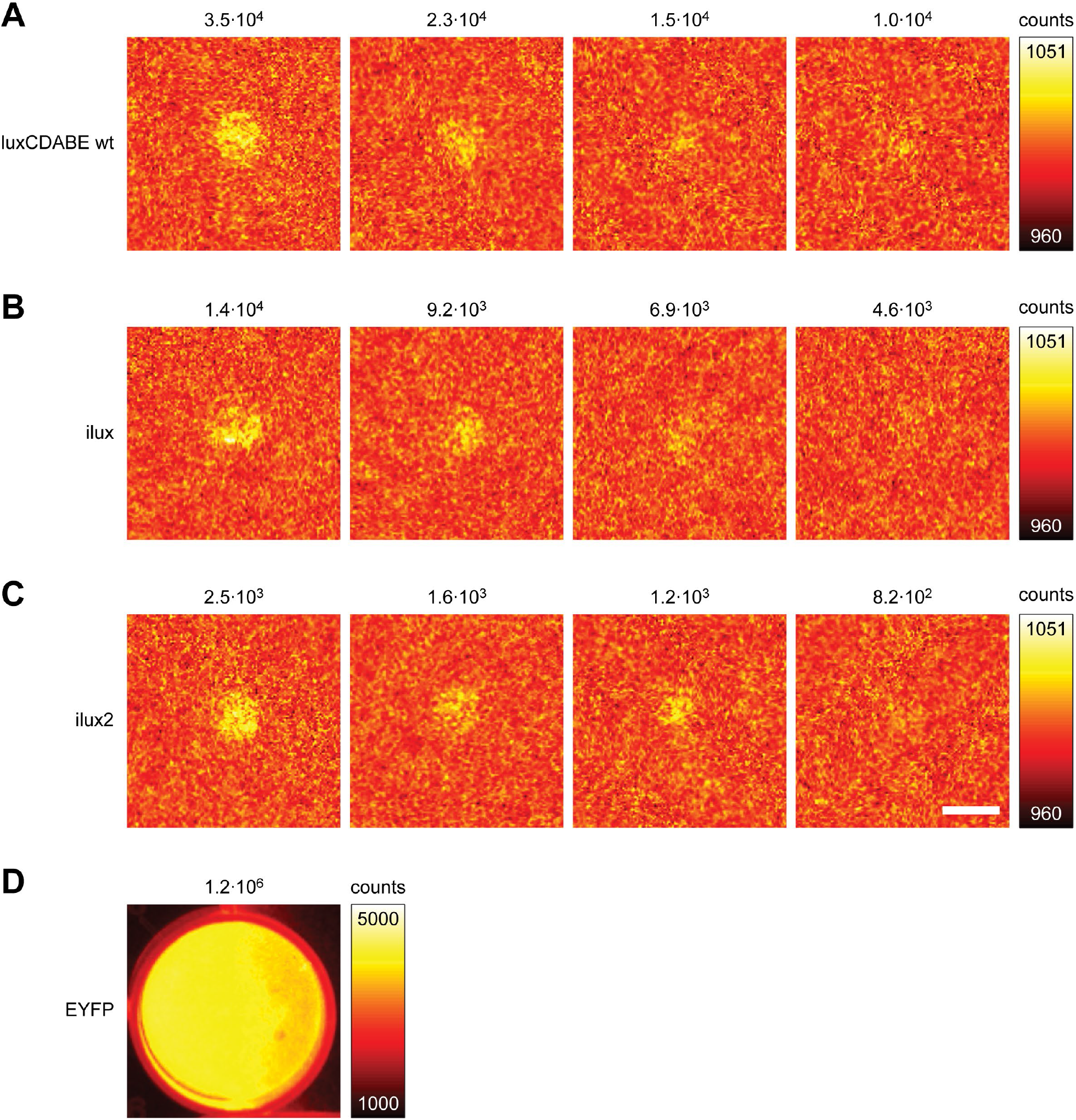
Detection of Top10 strains chromosomally labeled with (**A**) *luxCDABE wt*, (**B**) *ilux*, (**C**) *ilux2* or (**D**) EYFP on mashed potatoes. The indicated numbers of colony-forming units were applied in 1 µl PBS onto mashed potatoes in 24-well plates. Imaging was performed at room temperature. Bioluminescence from the *luxCDABE wt, ilux* and *ilux2* strains was imaged with an exposure time of 10 min. Fluorescence of the EYFP strain was excited at 520 nm and imaged with an exposure time of 1 s. Scalebar: 5 mm.

Imaging with the AI 600 RGB was performed at room temperature instead of 37 °C, where bioluminescence levels are 2–3 times higher ^16^. In addition, a relatively large distance between sample and objective lens was used to record a large field of view. Therefore, even lower cell numbers should be detectable under different imaging conditions. To test if even single cells can be detected, the *luxCDABE wt, ilux* and *ilux2* strains were imaged with a custom-built microscope as described in ^4, 16, 33^. As can be seen in Figure 10, only the *ilux2* strain could be imaged with a high signal-to-noise ratio, demonstrating the usefulness of this strain also for single-cell applications.

**Figure 10.**
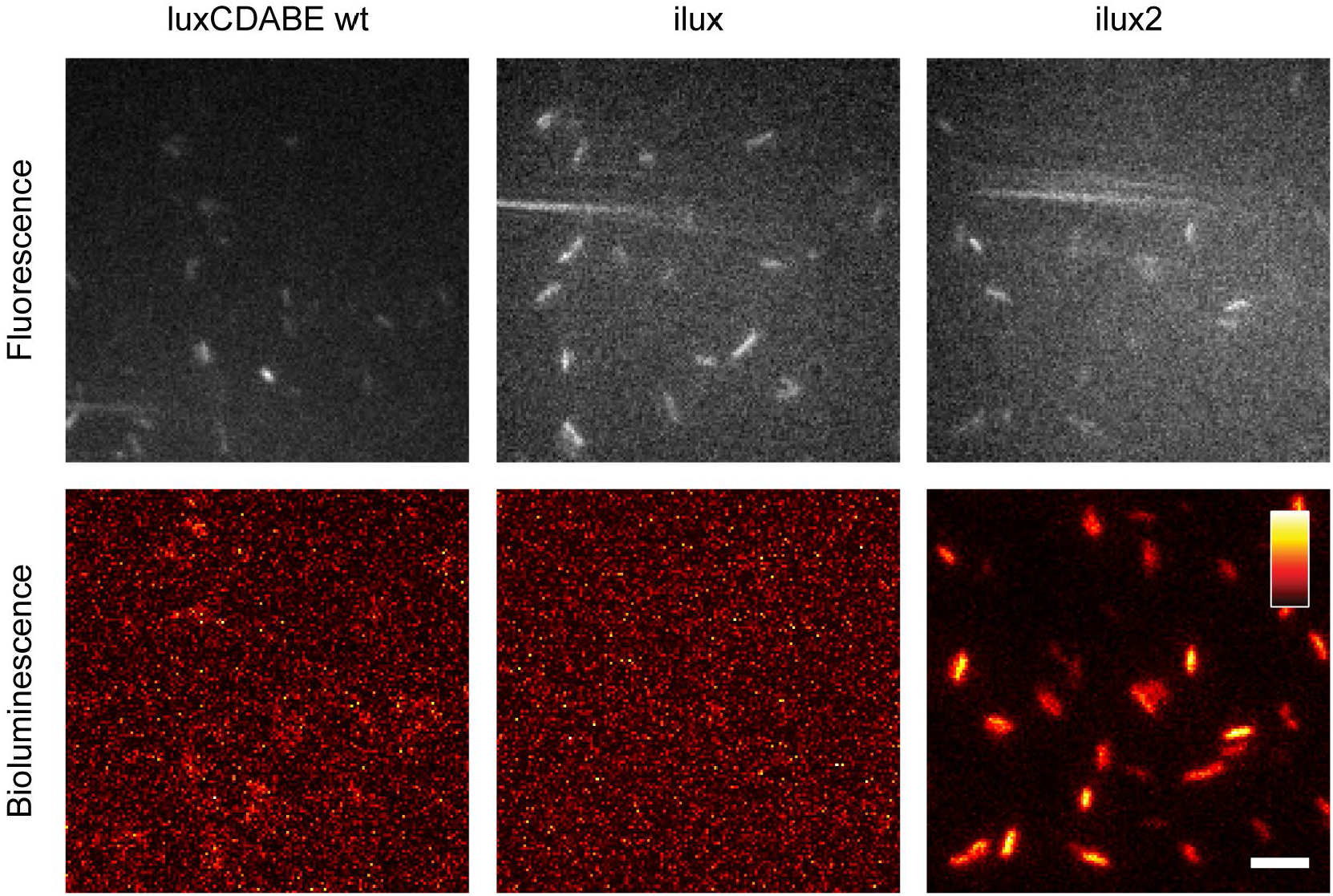
Imaging of single Top10 cells with *luxCDABE wt, ilux* or *ilux2* inserted into the chromosome. All three strains were transformed with EYFP pGEX(-) to detect the cells by their fluorescence excited at 491 nm (top row). Bioluminescence images were taken with an exposure time of 10 min. Imaging was performed at 37 °C. The colormap of each image was scaled to the minimum and maximum pixel values. Scalebar: 5 µm.

Finally, the *ilux2* strain was applied for long-term imaging of growth of *E. coli* cells in different food samples (Figure 11, Movies 1–5). Since only viable cells emit bioluminescence, both growth and spreading of the cells in the sample can be monitored over time. Cells grown on slices of raw cucumber and potato (Figure 11A,B, Movies 1 and 2) reached maximum bioluminescence after around 5 and 8 hours, respectively. Interestingly, the signal was not homogeneously distributed over the inoculated region, but especially on potato numerous bright spots appeared that changed their position over time. It is speculated that these bright spots emerge in regions of elevated supply with nutrient such as starch which might originate from breakdown of the potato cells. On mashed potatoes and egg yolk, the bioluminescence signal in the inoculated region was more uniform (Figure 11C,D, Movies 3 and 4). The signal strongly increased over time and reached its maximum after more than 24 h, indicating a high nutrient availability that supports long-term growth at high cell numbers. Since movement of the cells on the semi-solid surfaces was very limited, the signal finally started to decay at the center of the inoculated region where nutrients were depleted first whereas at the edges high bioluminescence was sustained for longer times. For comparison, Figure 11E and Movie 5 show the bioluminescence from milk that was chosen as a liquid sample. Although the sample was not moved after inoculation, the cells quickly spread over the complete dish, but were not evenly distributed in the sample during the entire 2 d observation time. Maximum bioluminescence was observed after ∼11 h and at the end of the 48 h observation time bioluminescence was barely detectable, presumably due to the depletion of nutrients and a drop in pH value. Together, these results demonstrate the utility of *ilux2* as a highly sensitive reporter for long-term imaging of bacteria in samples with high autofluorescence at an excellent signal-to-noise ratio.

**Figure 11.**
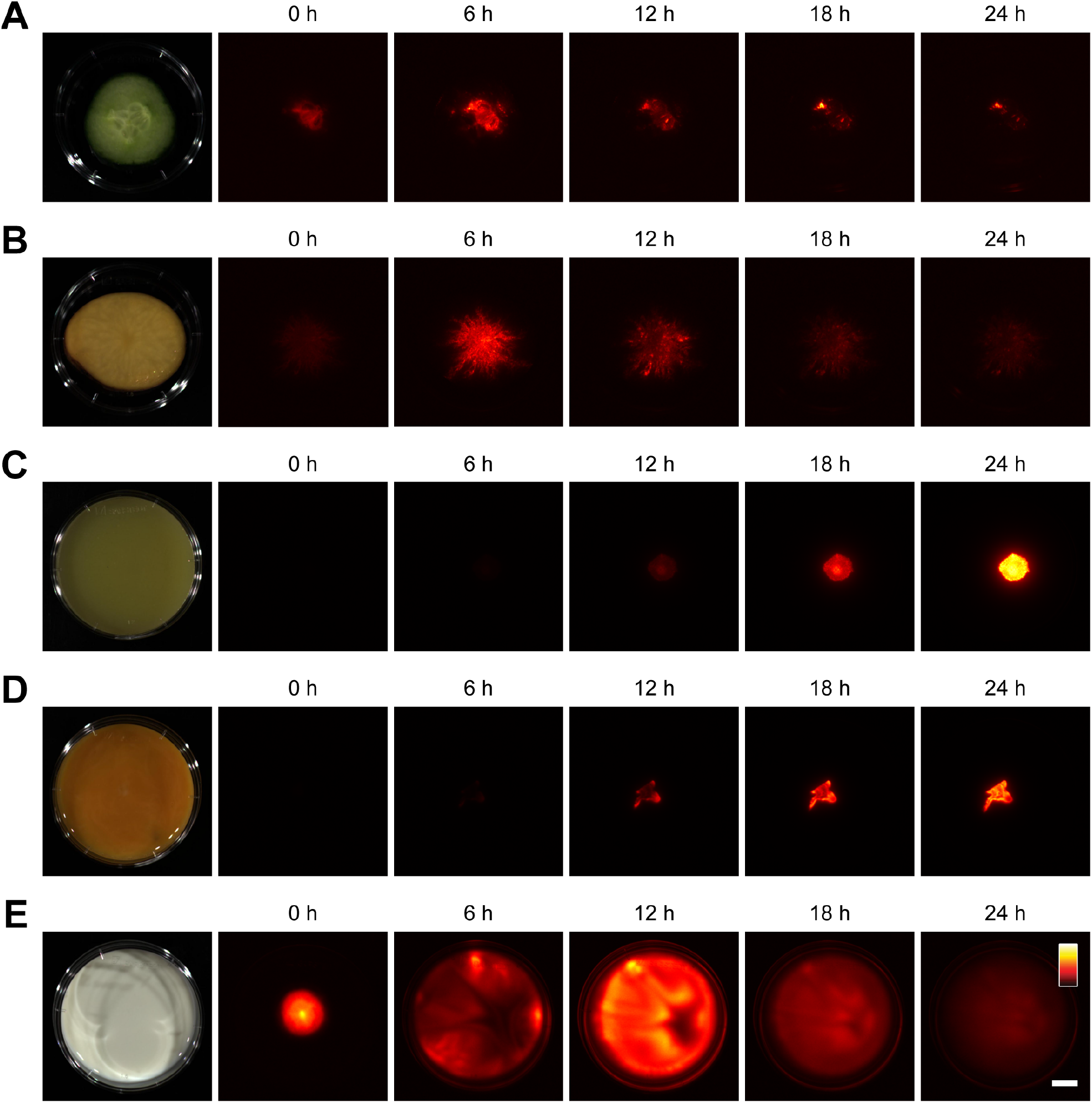
Growth of *E. coli* Top10 cells on different food products. Food samples were placed in 6 cm dishes and inoculated with Top10 cells chromosomally expressing *ilux2*. Bioluminescence images were taken at room temperature with an Amersham Imager AI 600 RGB after the indicated time periods. The colormap of each time series was scaled to the minimum and maximum pixel values. The following food products and exposure times per bioluminescence image were used: (**A**) cucumber, 1 min, (**B**) raw potato, 1 min, (**C**) mashed potatoes, 5 s, (**D**) egg yolk, 5 s, (**E**) milk, 1 min. Complete time series are shown in Movies 1–5. Scalebar: 1 cm.

## Discussion

By optimization of the screening conditions, it was demonstrated that light emission from the bacterial bioluminescence system can be improved under certain expression conditions compared to the previously engineered *ilux* system ^16^. This was achieved by component-wise mutagenesis of the involved proteins and screening under rate-limiting reaction conditions so that smaller improvements became detectable, which added up to a significant enhancement in light emission after multiple screening steps. It should be noted that no absolute value for the improvement in brightness can be specified since it strongly depends on the expression levels of all involved proteins. In addition, the overall process of bioluminescence is affected by the cellular availability of the substrates (e.g., fatty acids), resulting in a complex relationship between *lux* expression and bioluminescence emission. Moreover, toxic effects due to accumulation of the reaction products such as aldehydes can lead to adverse effects and reduce cell growth at elevated expression levels, which in turn decreases light emission. Therefore, bioluminescence from the newly generated *ilux2* operon is only increased at moderate expression levels. Nevertheless, the described screening system enables the generation of improved protein variants that would not be accessible otherwise. The presented screening strategy may therefore also be useful for the optimization of other processes involving multiple enzymes.

The enhanced light emission of the *ilux2* system results from 26 mutations in the fatty acid reductase complex (15 mutations in *iluxC*, 11 mutations in *iluxD*). The results demonstrate that the increased light output is at least in part due to the improved utilization of fatty acids other than tetradecanoic acid for which the fatty acid reductase complex has the highest activity ^34, 35^. A broadened substrate spectrum of iLux2CDE relative to LuxCDE wt would be in line with the high specificity of the wild-type fatty acid reductase complex for tetradecanoic acid since fatty acids that are potentially more abundant in the cell could also be used for the process of bioluminescence. The luciferase itself discriminates less between different chain lengths of its aldehyde substrate ^27, 28^ so that elevated production of aldehydes other than tetradecanoic aldehyde is expected to increase light emission. Besides the saturated fatty acids that were investigated in this study, also other substrates may be used with different specificity in the cell, such as unsaturated fatty acids or acyl-ACP and acyl-CoA esters.

The increased brightness of *ilux2* is particularly useful for the generation of stably autobioluminescent bacteria by chromosomal integration of the operon. These stable strains could for instance be used to image the spreading of bacterial infections in living animals and plants with enhanced sensitivity. Since bioluminescence emission requires metabolic activity, only living bacteria emit light. Therefore, bioluminescence imaging does not only yield information about the localization of the bacteria within the host, but also about their viability so that processes such as bacterial elimination by drugs or the immune system can be observed. In addition, *ilux2*-labeled strains can be used for *in vitro* applications, e.g. for drug screening to develop new antibacterial agents or to study antibiotic resistances. Another important application is the visualization of pathogenic bacteria on food samples to contribute to food safety. For example, bacterial growth can be studied under different storage conditions over long periods of time and different methods for preserving foodstuffs can be tested. It is therefore expected that *ilux2* will be a valuable reporter for highly sensitive bacterial imaging in samples that are not accessible by fluorescence measurements.

## Materials and Methods

### Calculation of optimal screening conditions

To determine the optimal conditions for the screening, a mathematical model was set up based on the reaction scheme in Figure 1. For better clarity, the following abbreviations are used below: A: aldehyde (RCHO), F: FMNH_2_, O: O_2_, L: LuxAB, LA: LuxAB·RCHO, LF: LuxAB·FMNH_2_, LFO: LuxAB·FMNHOOH, LFOA: LuxAB·FMNHOOR. It was first aimed at including also reactions for the production of aldehyde from fatty acids by LuxCDE and of FMNH_2_ from FMN by Frp in order to express the rate of light emission as a function of the reaction rate constants and concentrations of LuxAB, LuxCDE and Frp. However, since the resulting system was too complex to utilize its analytical solution, the concentrations of FMNH_2_ and aldehyde were used as variables instead of LuxCDE and Frp activity.

Based on the reaction scheme, the emitted bioluminescence B is proportional to the concentration of the LFOA complex:

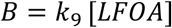

Under cellular steady-state conditions, the concentrations of LA, LF, LFO and LFOA remain constant. Therefore, the following rate equations can be deduced from Figure 1:

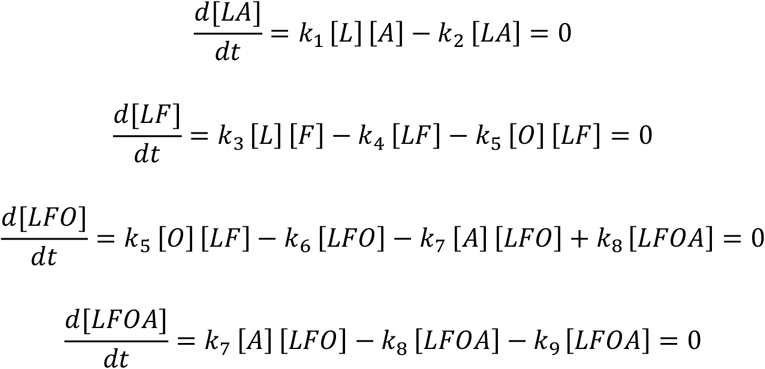

In addition, the concentrations of all intermediates must equal the total cellular concentration of luciferase [L]_0_ (for simplicity, other existing states of the luciferase are neglected since their concentration is also constant under steady-state conditions):

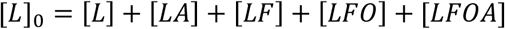

This set of six equations was solved with a custom-written Matlab (The MathWorks, Inc., Natick, Massachusetts) script. Since the rate constants k_i_ for the utilized proteins are unknown, they were all set to 1 for a qualitative description of the process, yielding the following equation for the bioluminescence emission:

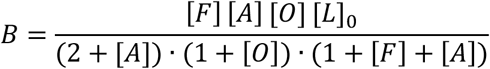

From this equation, it is evident that the bioluminescence signal increases less than proportionally with the substrate concentrations [F] and [A], i.e., an increase in [F] or [A] results in a smaller percentage change of B. For optimal sensitivity in the screening, the relative signal change with varying concentrations of the reactants (F, A and L, respectively) must be maximal. This was calculated as the derivative of B with respect to the screened reactant and normalized to B, yielding for F:

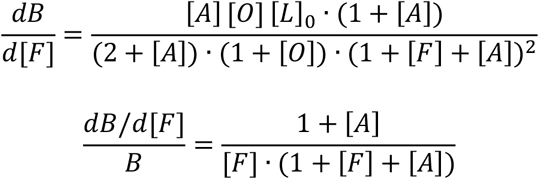

and for A:

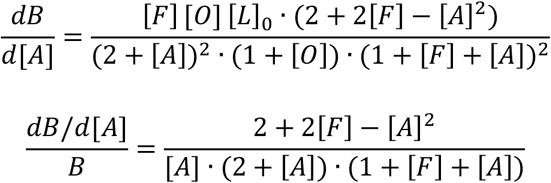

These results are plotted in Fig. 2 with [O] and [L]_0_ set to 1.

### Construction of screening vectors

The vector pGEX(-) for tagless expression of the Lux proteins was generated as described previously ^16^. For the two-plasmid based screening, another expression vector pGEX(-) Kan was generated in which the ampicillin resistance gene was exchanged against a kanamycin resistance marker in the following way: the kanamycin resistance gene was amplified by PCR using the primers KanR SpeI fwd and KanR AscI rev. The vector backbone was amplified with the primers pGEX(-) SpeI rev and pGEX(-) AscI fwd. Both parts were digested with SpeI and AscI and subsequently ligated.

For low-level expression, the tac promoter in pGEX(-) was exchanged against a lac promoter by PCR of the vector with the primers pGEX(-) +lac BamHI rev and pGEX(-) BamHI fwd and digestion with BamHI, followed by ligation without an insert.

To maintain two plasmids in the same cell during the screening, the ColE1 origin of replication in pGEX(-) was exchanged against the p15A origin of replication (p15Aori). For this purpose, the p15A origin of replication was PCR amplified with the primers p15Aori BglII fwd and p15Aori AvrII rev, whereas the vector backbone was amplified with the primers pGEX(-) BglII rev and pGEX(-) AvrII fwd. Both parts were digested with BglII and AvrII and subsequently ligated.

### Error-prone mutagenesis and screening

For mutagenesis of *luxAB, iluxAB* was amplified from *ilux* pGEX(-) described in ^16^ and cloned into the vector pGEX(-) lac p15Aori between the BamHI and NotI restriction sites. The remaining genes *iluxCDEfrp* were cloned into pGEX(-) Kan for high-level expression between the BamHI and NotI restriction sites. Error-prone mutagenesis of *iluxAB* in pGEX(-) lac p15Aori was performed as described previously ^16^. For screening, the ligation mixture was transformed into electrocompetent *E. coli* Top10 cells already containing the plasmid *iluxCDEfrp* pGEX(-) Kan. The cells were then spread out on LB agar plates containing ampicillin and kanamycin (both 50 µg/ml) and incubated overnight at 37 °C. Expression was not induced with IPTG. The following day, the plates were imaged with an Amersham Imager 600 RGB (AI 600 RGB, GE Healthcare, Chicago, Illinois) and the clones with the highest bioluminescence signal were selected and used for the next round of mutagenesis. Screening and mutagenesis of *luxCDE* were performed in analogy to *luxAB* using the plasmids *iluxCDE* pGEX(-) lac p15Aori and *iluxABfrp* pGEX(-) Kan.

For mutagenesis of frp, a Top10 *fre* knock-out (freKO) strain was generated in order to reduce FMNH_2_ production by the endogenous flavin reductase encoded by the *fre* gene. The knock-out strain was constructed by CRISPR using the system described in ^36^ with the target sequence GCGGCCTTTTCTTTTCGTGC and the oligonucleotide T*C*A*T*CCATCACTACCATCAAATACTGAGCGGCTCAAAAAGAAAAGGCCGCGTCTGGCACGATGCGGACACGATATACGGT for recombineering (* indicates phosphorothioate bonds).

### Comparison of LuxCDE wt, iLuxCDE and iLux2CDE activity with different fatty acids

*luxCDE wt, iluxCDE, ilux2CDE* and *iluxABfrp* were expressed in Top10 cells from the vector pGEX(-) Kan. For each construct, four colonies were spread out onto LB agar plates containing 50 µg/ml kanamycin and 20 µM IPTG and incubated overnight at 37 °C. The following day, cells were resuspended in LB medium and the OD_600_ was adjusted 0.5 (*luxCDE wt, iluxCDE, ilux2CDE*) or 1.0 (*iluxABfrp*). 100 µl of the *luxCDE* cell suspension was mixed with 3 µl of fatty acid solution (octanoic, decanoic, dodecanoic, tetradecanoic, hexadecanoic or octadecanoic acid, all dissolved in ethanol at a concentration of 10 mM) and incubated for 5 min at room temperature. Next, 10 µl of the *luxABfrp* cell suspension were added and incubated for 5 min at room temperature before starting the measurement. Bioluminescence intensity was determined with the AI 600 RGB. The backgound signal from a sample without addition of fatty acid was subtracted.

### *Lux* expression in *Pseudomonas fluorescens*

*Pseudomonas fluorescens* (DSM 50090) was obtained from DSMZ (Leibniz Institute DSMZ - German Collection of Microorganisms and Cell Cultures GmbH, Braunschweig, Germany) and cultivated in LB medium at 30 °C. For expression in *P. fluorescens, luxCDABE wt, ilux* and *ilux2* were inserted into the expression vector pJOE7771.1 ^31^ between the BamHI and SpeI restriction site. Plasmid transformation was performed by electroporation in 2 mm cuvettes at 2.5 kV using a capacitance of 25 µF and a resistance of 200 Ω. Cells were grown at 30 °C on LB agar plates containing 50 µg/ml kanamycin.

### Generation of stably bioluminescent *E. coli* strains

For chromosomal insertion, *luxCDABE wt, ilux, ilux2* and EYFP were first cloned into a pGEX(-) vector containing an AscI restriction site in front of the tac promoter. This vector was generated by PCR of pGEX(-) with the primers pGEX(-) AscI tac fwd and pGEX(-) AscI tac rev, followed by digestion of the PCR product with AscI and ligation without an insert. In the next step, *luxCDABE wt, ilux, ilux2* and EYFP were cut from previously generated pGEX(-) plasmids with BamHI and NotI and ligated into the new vector pGEX(-) AscI tac digested with the same enzymes. Finally, the complete inserts including the tac promoter were cut out from the newly generated plasmids using AscI and NotI and ligated into the vector pGRG25 described in ^32^ that was digested with the same enzymes.

Insertion of the transgenes from pGRG25 into *E. coli* Top10 was performed as described in ^32^. Curing of the pGRG25 plasmid was checked by ampicillin sensitivity. Correct insertion of the transgenes into the chromosome was verified by colony PCR of the final strains using the primers attTn7 fwd and luxC EcoRI rev for *luxCDABE wt, ilux* and *ilux2*, frp SalI fwd and attTn7 rev for *ilux* and *ilux2* and attTn7 fwd and attTn7 rev for EYFP.

### Bioluminescence imaging

To compare the brightness of different variants, bacteria were transformed with the indicated plasmids and grown on LB agar plates containing the appropriate antibiotics overnight. The following day, single colonies were spread out onto a new LB agar plate. If not stated otherwise, plates were incubated at 37 °C overnight. The following day, plates were imaged with the AI 600 RGB to quantify the bioluminescence signal.

The number of detectable cells of stable Top10 strains containing *luxCDABE wt, ilux, ilux2* and EYFP on mashed potatoes was compared in 24-well plates. Each well was filled with freshly prepared mashed potatoes (consisting of boiled potatoes and water only) up to a height of around 0.5 cm. Cells grown at 37 °C on LB agar plates without antibiotics or IPTG were resuspended in PBS and 1 µl of different dilutions of the cell suspension was added at the center of each well. Plates were imaged at room temperature with the AI 600 RGB. For the *luxCDABE wt, ilux* and *ilux2* strains, bioluminescence was recorded for 10 min. For the EYFP strain, fluorescence was recorded with an excitation wavelength of 520 nm, a 605BP40 filter and a 1-s exposure time. The number of viable cells was determined by plating different dilutions of the cell suspensions on LB agar plates and counting the number of colony-forming units (cfu) after incubation overnight at 37 °C.

Imaging of single *E. coli* cells was performed using a custom-built microscope as described previously ^4, 16, 33^. Briefly, cells grown on LB agar plates were resuspended in LB medium and imaged between a coverslip and LB agar pad. Imaging was performed in a stage top incubation system (Okolab, Ottaviano, Italy) at 37 °C. A 60× oil immersion objective lens (PlanApo N 60x/1.42, Olympus, Shinjuku, Tokyo, Japan) and an electron-multiplying charge-coupled device (EMCCD) camera (iXon EMCCD DU-897E-CS0-#BV, Andor Technology Ltd, Belfast, Northern Ireland) with the camera sensor cooled to –100 °C were used to detect the bioluminescence light. To select and focus the cells, cells were transformed with EYFP pGEX(-) beforehand. EYFP fluorescence was excited with a laser (Cobolt Calypso 50 mW, Hübner Photonics, Kassel, Germany) at 491 nm. Bright pixels were filtered out using a custom-written Matlab script as described in ^4, 16, 33^.

Movies of the stable *ilux2* Top10 strain on food products were taken with the AI 600 RGB. Slices of raw cucumber and potato, raw egg yolk, mashed potatoes (consisting of boiled potatoes and water only) and UHT milk were placed in transparent 6-cm dishes, inoculated with around 5·10^6^ cfu in PBS and covered with a lid to prevent dehydration of the samples. One image (exposure time 1 min or 5 s) was recorded per hour for 48 h.

## Acknowledgements

I wish to thank Stefan W. Hell for providing access to the optical setup for single-cell imaging and Steffen J. Sahl for feedback on the manuscript. pKDsgRNA-ack, pCas9-CR4 and pKDsgRNA-p15 were a gift from Kristala Prather (Addgene plasmids #62654–62656). pGRG25 was a gift from Nancy Craig (Addgene plasmid #16665). pJOE7771.1 was a gift from Josef Altenbuchner (Addgene plasmid #135077).

## Movies

**Movie 1**

*E. coli* Top10 strain chromosomally expressing *ilux2* grown on cucumber. A slice of cucumber was inoculated with 1·10^6^ colony-forming units (cfu). Bioluminescence images were taken at room temperature with an Amersham Imager AI 600 RGB using an exposure time of 1 min per image.

**Movie 2**

*E. coli* Top10 strain chromosomally expressing *ilux2* grown on potato. A slice of raw potato was inoculated with 1·10^6^ cfu. Bioluminescence images were taken at room temperature with an Amersham Imager AI 600 RGB using an exposure time of 1 min per image.

**Movie 3**

*E. coli* Top10 strain chromosomally expressing *ilux2* grown on mashed potatoes. A dish filled with mashed potatoes was inoculated with 5·10^5^ cfu. Bioluminescence images were taken at room temperature with an Amersham Imager AI 600 RGB using an exposure time of 5 s per image.

**Movie 4**

*E. coli* Top10 strain chromosomally expressing *ilux2* grown on egg yolk. A dish filled with raw egg yolk was inoculated with 5·10^5^ cfu. Bioluminescence images were taken at room temperature with an Amersham Imager AI 600 RGB using an exposure time of 5 s per image.

**Movie 5**

*E. coli* Top10 strain chromosomally expressing *ilux2* grown in milk. A dish filled with UHT milk was inoculated with 3·10^6^ cfu. Bioluminescence images were taken at room temperature with an Amersham Imager AI 600 RGB using an exposure time of 1 min per image.

## Additional Files

**Supplementary file 1**

Nucleotide sequences of the *ilux2* genes. The sequences of *ilux2A, ilux2B* and *ilux2frp* are identical to *iluxA, iluxB* and *iluxfrp*, respectively. The sequence of *ilux2E* is the same as in the wild-type *lux* operon.

