## Supplementary File 1 for "Generation of bright autobioluminescent bacteria by chromosomal integration of the improved *lux* operon *ilux2*"

>i*lux2C*

ATGACTAAAAAAATTTCATTCATTATTAATGGCCAGGTAGAAATCTTTCCCGAAAGCGATGAT
TTAGTGCAATCCATCACTTTTGGTGATAATAGTGTTTACCTGCCAATATTGAACGACTCTCAT
GTAAAAACCATTACTGATTGTAATGGAAATAACGACTTACGGTTGCATGACATTGTCAATTTT
CTCTATACGGTAGGGCAAAGATGGAAAAATAATGAATACCCAAGACGCAGGACATACATTC
GTGACTTAAAAAAATATATGGGATATTCAGAAGAAATGGCTAAGCTAGAGGCCAATTGGATA
TCTATGATATTATGTTCTAAAGGCGGCCTTTATGATGTTGTAGAAAATGAACTTGGTTCTCGC
CATATCATGGATGAATGGATACCTCAGGGTGAAAGTTATGTTCGGGCTTTTCCGAAAGGTAA
ATCCGTACATCTGTTGGCAGGTAATGTTCCATTATCTGGGATCATGTCTATATTACGTGCAA
TTTTAACTAAGAATCAGTGTATTATAAAAACATCGTCAACCGATCCTTTTACCGCTAATGCAT
TAGCTTTAAGTTTTATTGATGTAGACCCTAATCATCCGATAACGCGCTCTTTATCTGTTATAT
ATTGGCCCCACCAAGGTGATATATCACTCGCAAAAGAAATTATGCGACATGCGGATGTTATT
GTCGCATGGGGAGGGCCAGATGCGATTAATTGGGCGGTAGAGCATGCGCCATCTAATGCT
GATGTGATTAAATTTGGTCCTAAAAAGAGCCTTTGCATTATCGATAATCCTGTTGATCTGAC
GTCTGCAGCGACAGGTGCGGCTCATGATGTTTGTTTTTACGATCAGCGAGCTTGCTTTTCT
GCCCAAAACATATATTATATGGGAAATCATTATGAGGAATTTAAGTTGGCATTGATAGAAAAA
CTTAATCTATATGCGCATATATTACCGAATGCAAAAAAAGATTTTGATGAAAAGGCGGCCTA
TTCTTTAGTTCAAAAAGAGAGCTTGTTTGCCGGATTAAATGTAGAGGCGGATATTCATCAAC
GTTGGACGATTATTGAGTCAGATGCAGGTGTGGAATTTAATCAACCACTTGGCAGATGTGT
GTACCTTCATCACGTCGATAATATTGAGCAAATATTGCCTTATGTTCAAAAAAATAAGACGCA
AACCATATCTATTTTTCCTTGGGAGTCATCATTTAAATATCGAGATGCGTTAGCATTAGAAGG
TGCGGAAAGGATTGTAGAAGCAGGAATGAATAACATATTTCGAGTTGGTGGATCTCATGAC
GGAATGAGACCGTTGCAACGATTAGTGAAGTATATTTCTCATGAAAGGCCATCTAACTATAC
GGCTAAGGATGTTGCGGTTGAAATAGAACAGACTCGTTTCCTGGAAGAAGACAAGTTCCTT
GTATTTGTCCCATAA

>i*lux2D*

ATGGAAAAAGAATCAAAATATAAAACCATCGACCACGTTATTTGTGTTGAAGGAAATAAAAAA
ATTCATGTTTGGGAAACGCTGCCAGAAGAAAACAGCCCAAAGAGAAAGAATGCCATTATTAT
TGCGTCTGGTTTTGCCCGCAGGATGGATCATTTTGCTGGACTGGCGGAATATTTATCGCGG
AATGGGTTTCATGTGATCCGCTATGATTCGCTTCACCACGTTGGATTGAGTTCAGGGACAAT
TGATGAATTTACAATGTCTACAGGAAAGCAGAGCTTGTTAGCAGTGGTTGATTGGTTAACTA
CACGAAATATAAGTAACTTCGGTGTGTTGGCTTCAAGCTTATCTGCGCGGATAGCTTATGCA
AGCCTATCTGAAATCAATGCTTCGTTTTTAATCACCGCAGTCGGTGTTGTTAACTTAAGATAT
TCTCTTGAAAGAGCTTTAGGGTTTGATTATCTCAGTCTTCCCATTAATGAATTGCCGAAAAAT
CTAGTTTTTGAAGGCCATAAATTGGGTGCTGAAGTCTTCGCGAGAGATTGTCTTGATTTTGG
TTGGGAAGACTTAGCTTCTACAATTAATAACATGATGTATCTTGATATACCGTTTATTGCTTT
TACCGCAAATAACGACAATTGGGTCAAGCAAGATGAAGTTATCACATTGTTATCAAATATTC
GAAGTAATCGATGTAGGATATATTCTTTGTTAGGAAGTTCGCATGACTTGAGTGAAAATTTA
GTGGCCCTGCGCAATTTTTATCAATCGGTTACGAAAGCCGCTATCGCGATGGATAATGATC
ATCTGGATATTAATGTTGATATTACTGAACCGTCATTTGAACATTTAACTATTGCGACAGTCA
ATGAACGCCGAATGAGAATTGAGATTGTAAATCAAGCAATTTCTCTGTCTTAA

>i*lux2A*

ATGAAATTTGGAAACTTTTTACTTACATACCAACCTCCCCAATTTTCTCAAACAGAGGTAATG
GAACGTTTGGTTAAATTAGGTCGCATCTCTGAGGAGTGTGGTTTTGATACCGTATGGTTACT
GGAGCATCATTTCACGGAGTTTGGTCTACTTGGTAACCCTTATGTCGCTGCTGCATATTTAC
TTGGCGCGACTAAAAAATTGAATGTAGGAACCGCCGCTATTGTTCTTCCCACAGCCCATCC
AGTACGCCAACTTGAAGATGTGAATTTATTGGATCAAATGTCAAAAGGACGATTTCGGTTTG
GTATTTGCCGAGGGCTTTACAACAAGGACTTTCGCGTATTCGGCGCGGATATGAATAACAG
TCGCGCCTTAGCGGAATGCTGGTACGGGCTGATAAAGAATGGCATGACAGAGGGATATAT
GGAAGCTGATAATGAACATATCAAGTTCCATAAGGTAAAAGTAAACCCCGCGGCGTATAGC
AGAGGTGGCGCACCGGTTTATGTGGTGGCTGAATCAGCTGCGACGACTGAGTGGGCAGCT
CAATTTGGCCTACCGATGATATTAAGTTGGATTATAAATACTAACGAAAAGAAAGCACAACTT
GAGCTTTATAATGAGGTGGCTCAAGAATATGGGCACGATATTCATAATATCGACCATTGCTT
ATCATATATAACATCTGTAGATCATGACTCAATTAAAGCGAAAGAGATTTGCCGGAAATTTCT
GGGGCATTGGTATGATTCTTATGTGAATGCTACGACTATTTTTGATGATTCAGACCAAACAA
GAGGTTATGATTTCAATAAAGGGCAGTGGCGTGACTTTGTATTAAAAGGACATAAAGATACT
AATCGCCGTATTGATTACAGTTACGAAATCAATCCCGTGGGAACGCCGCAGGAATGTATTG
ACATAATTCAAAAAGACATTGATGCTACAGGAATATCAAATATTTGTTGTGGATTTGAAGCTA
ATGGAACAGTAGACGAAATTATTGCTTCCATGAAGCTCTTCCAGTCTGATGTCATGCCATTT
CTTAAAGAAAAACAACGTTCGCTATTATATTAG

>i*lux2B*

ATGAAATTTGGATTGTTCTTCCTTAACTTCATCAATCCGACAACTGTTCAAGAACAAAGTATA
GTTCGCATGCAGGAAATAACGGAGTATGTTGATAAGTTGAATTTTGAACAGATTTTAGTGTA
TGAAAATCATTTTTCAGATAATGGTGTTGTCGGCGCTCCTCTGACTGTTTCTGGTTTTCTGCT
CGGTTTAACAGAGAAAATTAAAATTGGTTCATTAAATCACATCATTACAACTCATCATCCTGT
CCGCATAGCGGAGGAAGCTTGCTTATTGGATCAGTTAAGTGAAGGGAGATTTATTTTAGGG
TTTAGTGATTGCGAAAAAAAAGATGAAATGCATTTTTTTAATCGCCCGGCTGAATATCAACA
GCAACTATTTGAAGAGTGTTATGAAATCATTAACGATGCTTTAACAACAGGCTATTGTAATCC
AGATAACGATTTTTATAGCTTCCCTAAAATATCTGTAAATCCCCATGCTTATACGCCAGGCG
GACCTCGGAAATATGTAACAGCAACCAGTCATCATATTGTTGAGTGGGCGGCCAAAAAAGG
TATTCCTCTCATCTTTAAGTGGGATGATTCTAATGATGTTAGATATGAATATGCTGAAAGATA
TAAAGCCGTCGCGGATAAATATGACGTTGACCTATCAGAGATAGACCATCAGTTAATGATAT
TAGTTAACTATAACGAAGATAGTAATAAAGCTAAACAAGAGACGCGTGCATTTATTAGTGATT
ATGTTCTTGAAATGCACCCTAATGAAGATTTCGAAAATAAACTTGAAGAAATAATTGCAGAAA
ACGCTGTCGGAAATTATACGGAGTGTATAACTGCGGCTAAATTGGCAATTGAAAAGTGTGG
TGCGAAAAGTGTATTGCTGTCCTTTGAACCAATGAATGATTTGATGAGCCAAAAAAATGTAA
TCAATATTGTTGATGATAATATTAAGAAGTACCACATGGAATATACCTAA

>i*lux2E*

ATGACTTCATATGTTGATAAACAAGAAATTACAGCAAGCTCAGAAATTGATGATTTGATTTTT
TCGAGCGATCCATTAGTGTGGTCTTACGACGAGCAGGAAAAAATCAGAAAGAAACTTGTGC
TTGATGCATTTCGTAATCATTATAAACATTGTCGAGAATATCGTCACTACTGTCAGGCACACA
AAGTAGATGACAATATTACGGAAATTGATGACATACCTGTATTCCCAACATCGGTTTTTAAGT
TTACTCGCTTATTAACTTCTCAGGAAAACGAGATTGAAAGTTGGTTTACCAGTAGCGGCACG
AATGGTTTAAAAAGTCAGGTGGCGCGTGACAGATTAAGTATTGAGAGACTCTTAGGCTCTG
TGAGTTATGGCATGAAATATGTTGGTAGTTGGTTTGATCATCAAATAGAATTAGTCAATTTGG
GACCAGATAGATTTAATGCTCATAATATTTGGTTTAAATATGTTATGAGTTTGGTGGAATTGT
TATATCCTACGACATTTACCGTAACAGAAGAACGAATAGATTTTGTTAAAACATTGAATAGTC
TTGAACGAATAAAAAATCAAGGGAAAGATCTTTGTCTTATTGGTTCGCCATACTTTATTTATT
TACTCTGCCATTATATGAAAGATAAAAAAATCTCATTTTCTGGAGATAAAAGCCTTTATATCA
TAACCGGAGGCGGCTGGAAAAGTTACGAAAAAGAATCTCTGAAACGTGATGATTTCAATCA
TCTTTTATTTGATACTTTCAATCTCAGTGATATTAGTCAGATCCGAGATATATTTAATCAAGTT
GAACTCAACACTTGTTTCTTTGAGGATGAAATGCAGCGTAAACATGTTCCGCCGTGGGTATA
TGCGCGAGCGCTTGATCCTGAAACGTTGAAACCTGTACCTGATGGAACGCCGGGGTTGAT
GAGTTATATGGATGCGTCAGCAACCAGTTATCCAGCATTTATTGTTACCGATGATGTCGGGA
TAATTAGCAGAGAATATGGTAAGTATCCCGGCGTGCTCGTTGAAATTTTACGTCGCGTCAAT
ACGAGGACGCAGAAAGGGTGTGCTTTAAGCTTAACCGAAGCGTTTGATAGTTGA

>i*lux2frp*

ATGGTGAAGATACAGCCCATCCCCACAACTAGCCAGGGCAGCCTTTTTATAATGAATAGCA
CCATAGAGACAATCCTGGGCCATAGATCCATTAGGAAGTTCACATCTGAACCTATTGCTAGT
GAGCAGCTGCAAACGATTCTTCAGTCTGGGCTCGCTGCTTCAAGCTCATCCATGCTGCAGG
TTGTGAGTATAATTCGGGTTACAGACACGGAAAAGAGAAAATTGCTCGCTCAATATGCCGG
CAACCAGACGTATGTGGAATCCGCTGCTGAGTTCCTGGTCTTTTGTATAGACTACCAGCGA
CACGCTACTATCAACCCCGATGTCCAAGCTGACTTTACCGAGCTGACCCTGATTGGTGCAG
TGGATTCCGGCATAATGGCCCAGAATTGCCTCCTGGCAGCAGAATCAATGGGTCTTGGCG
GAGTCTATATCGGAGGACTTCGGAACTCAGCTGCCCAAGTGGATGAGTTGCTCGGACTGC
CCAAGAACACAGCTATCCTCTTCGGAATGTGCTTGGGGCACCCCGATCAGAGCCCTGAGA
CAAAGCCTAGACTGCCCGCTCATGTGATCGTGCACGAGAACCAATATCAAGCTCTGAACAT
TGACGACGTACAGGCGTATGACAAAACATTGCAGGAGTATTATGCCAGCAGAACCAGCAAC
CAGAAGCAGAGTGTCTGGTCCCAGGAAACTGCAGGCAAGCTGGCCGGAGAATCCCTCCCA
CACATCCTGCCATACCTGAACTCCAAGGGCCTTGCCAGAAGGTAA
